# Multi-view gene panel characterization for spatially resolved omics

**DOI:** 10.1101/2025.02.19.639174

**Authors:** Daniel Kim, Wenze Ding, Akira Nguyen Shaw, Marni Torkel, Cameron J Turtle, Pengyi Yang, Jean Yang

## Abstract

Spatially resolved transcriptomics has transformed our ability to study complex tissues at the cellular and subcellular resolution. However, targeted spatial technologies require pre-selected gene panels, which are typically curated based on existing biological knowledge and prior research hypotheses. While current methods often prioritize capturing cell type information, we argue that an effective gene panel should also capture cell type diversity, cell states, pathway-level information, and minimize redundancy. To address these broader requirements, we developed a gene panel characterization platform that characterizes panels across multiple perspectives, thus allowing us to compare panels comprehensively. Notably, computationally constructed gene panels performed competitively in capturing major cell types when compared to our in-house manually curated panel. However, refined manual curation offered distinct advantages, particularly in capturing minor and rare cell types and exhibited lower information redundancy comparatively. Building on this framework, we integrated these metrics into a deep learning platform, panelScope, leveraging them as a loss function to design holistic gene panels. Using an acute myeloid leukemia (AML) dataset with 42 well-defined cell types and the 5K Xenium panel from 10X Genomics, we demonstrate the utility of our framework in comprehensively characterizing gene panels, enabling the design of tailored panels that address diverse research needs.

## 1. Introduction

The emergence of spatially resolved transcriptomics (SRT) has transformed our ability to quantify gene expression *in situ* while preserving the native tissue environment. This technological leap has revealed which genes are expressed and where they are expressed within complex tissues. This spatial context is critical for elucidating cell–cell interactions, tissue heterogeneity, and the architectural nuances that govern disease progression and therapeutic response. Consequently, SRT has enabled insights that would be difficult or impossible to achieve using conventional sequencing alone (Garcia-Alonso *et al*., 2021; McNamara *et al*., 2021). The ability of SRT to capture spatial features of gene expression holds promise for advancing our understanding of tissue development, disorders, and other pathologies that require spatial organization of cell-types, cell-states, and transcripts.

There are a variety of SRT techniques, including fluorescence *in situ* hybridization (FISH)-based methods, such as seqFISH+ and MERFISH, which directly label and visualize transcripts within tissue sections (Chen *et al*., 2015; Eng *et al*., 2019). In contrast, spot-level RNA-sequencing techniques, such as SLIDE-seq and high-definition spatial transcriptomics (HDST), utilize barcoded bead arrays to capture RNA while retaining spatial coordinates (Rodriques *et al*., 2019). These SRT platforms differ significantly in their capabilities: sequencing-based approaches (e.g., SLIDE-seq, HDST) can capture whole-transcriptome data (un-targeted transcriptomics), while *in situ* hybridization methods generally focus on a targeted subset of genes of interest (targeted transcriptomics) (He *et al*., 2022). Targeted spatial transcriptomics is often employed when researchers wish to focus on a specific set of genes and do not require whole-transcriptome data. By sequencing a smaller subset of genes, they can achieve higher sensitivity and accuracy, which is particularly valuable for detecting lowly expressed genes. However, targeted spatial transcriptomics requires the careful design of a gene panel tailored to specific research objectives.

To facilitate panel design, several computational methods have been developed for designing gene panels. These methods can be broadly categorized into two categories: imputation-based methods and classification-based methods. The first category of imputation approaches select highly variable genes and assesses their contribution to the overall gene panel (Subramanian *et al*., 2017; Liang *et al*., 2021; Missarova *et al*., 2021; Kuemmerle *et al*., 2022). Examples in this category include geneBasis which selects for genes that preserve the original transcriptional variation in a reference dataset. They achieve this by computing the distance between a “true” k-NN representation of the full reference data and a k-NN representation using random genes (Missarova *et al*., 2021). Genes with the largest distances are then selected for the panel. Classification-based methods; include approaches that focus on selecting genes that can differentiate between cell types, emphasizing cell type-specific markers (Vargo and Gilbert, 2020; Aevermann *et al*., 2021; Dumitrascu *et al*., 2021). Examples in this category include gpsFISH, which uses a genetic algorithm to select genes that contribute to higher accuracy in cell type classification via cross validation (Zhang *et al*., 2024); and panelScope. To date, all these computational methods focus on optimizing panel characteristics like classification accuracy, but have overlooked other important criteria, such as minimising redundancy or capturing pathway-level information. A notable exception is Spapros, which incorporates additional criteria, beyond cell type classification accuracy, into its panel design process (Kuemmerle *et al*., 2022).

Although various panel design methods have been proposed, limited effort has been directed toward comprehensively characterizing gene panels. The desirable characteristics of a gene panel often depend on one’s research objectives. For instance, a panel designed to capture diverse pathways would be suitable for studying multiple biological processes with higher pathway diversity, while a panel focused on a specific disease might prioritize low pathway diversity to ensure thorough coverage of relevant processes. Currently, researchers are often left to manually assess the suitability of a selected gene panel, making it time consuming and challenging to align panel design with specific research goals. This highlights the need for a comprehensive framework for gene panel characterization, which is currently lacking.

To this end, we present panelScope, a framework based on a diverse collection of metrics to characterize a gene panel, allowing researchers to determine whether a panel is well-suited to their study’s objectives. We demonstrate the utility of panelScope by generating multiple-views of gene panels that describe their ability to capture cell types of interest, enrichment for biological pathways, or the amount of redundant information. In parallel, we leverage these metrics as loss functions in a genetic algorithm for panel design, where users can choose to weight each characterization category. Importantly, we have implemented this framework in an interactive web platform, which includes a library of pre-existing gene panels that users can compare to their own gene panels. Thus, by quantitatively summarizing a panel from multiple views, panelScope enables the design of panels that can capture diverse information relevant to one’s specific research questions.

## 2. Materials and Methods

### 2.1 Datasets

#### Dataset 1 - AML dataset

We illustrate our panelScope framework using a scRNA-seq dataset of bone marrow aspirates from 19 AML patients and a healthy individual (Beneyto-Calabuig *et al*., 2023). Both healthy hematopoietic differentiation and diseased myeloid states are represented in this dataset. We preprocessed the data by removing any genes with less than a total of 10 counts, cells with less than a total of 200 counts, and cell types with less than 10 cells. The final dataset included 39,146 genes, 3,437 cells, and 37 cell types, as annotated by the original authors.

#### Dataset 2 - Lymph node dataset

To demonstrate the utility of our spatial metrics we used the 5k Xenium Panel of human lymph nodes by 10 Genomics, available at: https://www.10xgenomics.com/resources/datasets (Accessed October 10, 2024). This dataset includes 4,624 genes, 708,983 cells, and 47,977,580 regions. Given the dataset’s size, we downsampled the original data to include 70,487 cells while retaining all 4,624 genes without applying any additional filtering.

### 2.2 Gene panel curation

We curated eight gene panels to apply our framework and illustrate how different panel properties can be characterized. Our first example uses the AML dataset and features eight gene panels generated through various strategies, including expert curation, random selection, computational methods, and the use of large language models.

*Random* - we selected a random set of genes (n=160) from the dataset.

*SEG* - a set of stably expressed genes, previously identified (Lin *et al*., 2019), (n=160). We selected the top n genes according to the stability scores provided in the original study.

*Expert-curated*- an in‐house‐designed AML gene panel (n=160). Potential markers for each cluster were identified by factorizing the expression matrix using non-negative matrix factorization. Markers were selected based on the specificity for each cluster, as verified by visual inspection of gene expression across dimension reduction plots.

*IO* - This is the Immune Oncology gene panel designed by 10X Genomics available from https://www.10xgenomics.com/products/xenium-panels (n=380).

*geneBasis* - panel selection using geneBasis after its initial gene screening function “retain_informative_genes”, all parameters were set to default values (n=117).

*gpsFISH* - we conducted panel selection by strictly following *gpsFISH*’s vignette (n=117). panelScope - optimization algorithm based on our developed metrics (see Section 2.X for details) (n = 200)

*chatGPT o1* - To explore the capabilities of *ChatGPT o1*, we crafted two distinct prompts asking it to design two separate AML‐specific gene panels (Supplementary 1). Each prompt instructed *ChatGPT o1* to maximize the characteristics set forth in our framework—namely, feature specificity, feature diversity, and pathway coverage. Because we wanted to include cell‐type‐specific markers, the prompts differed in how much cell‐type information was provided. Prompt 2 specified the cell types expected in AML (*ChatGPT o1*), while prompt 2 listed all 42 cell types present in the AML dataset (*ChatGPT o1+*). Both panels contained 200 genes each.

Our second example uses the lymph node dataset described above to demonstrate the spatial metrics of panelScope. We used the same protocol as described above, excluding the *Expert-curated* panel, to curate seven gene panels for this dataset. The two prompts used for this dataset can be found in the supplementary material (Supplementary 2).

### 2.3 panelScope characterization metrics

#### 2.3.1 Feature specificity

This category aims to assess a panel’s ability to distinguish groups of interest (e.g., cell types or cell states). We evaluate the performance of a random forest model trained on a dataset subsetted by a given panel’s genes. If the genes in a given panel are informative, we expect the model to perform well in identifying these groups. We quantify this using balanced accuracy and accuracy stratified by the groups of interest as metrics.

Additionally, we assess the cell type specificity of a gene panel of size *G*, which provides a measure of how uniquely a gene is expressed in a cell type relative to all other cell types. Suppose we aim to capture the set of all cell types *C* in our panel design, a natural uniqueness score is the measure the abundance of cells within a given cell type *c*∈*C* that express the gene relative to all other cell types *c*’ ∈*C*.

##### Gene specificity score (*gsc*)

This score aims to quantify the specificity of an individual gene for a given cell type. For a typical gene *g*, let us denote *x*_*cj*_ as the expression for the *jth* cell in cell type *c*∈*C, n*_*c*_ and *n*_*c’*_ represents the total number of cells in cell type *c*and all other cell types *c*’ respectively. We define the gene specificity score for gene *g* in cell type *c* as:

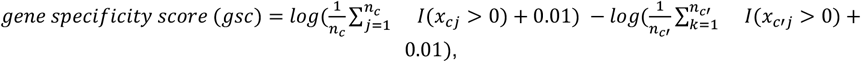

where *I*(*x>*0)is an indicator function and equals 1 if *x>*0, and 0 otherwise. Note that we typically have an arbitrary small value of 0.01 inside the logarithm to avoid taking the log of zero. A large specificity score indicates that gene *g* is a unique marker for cell type *c*, with a higher proportion of cells expressing the gene of interest in cell type *c*relative to all other cell types *c*’. In contrast, a negative specificity score suggests that the gene is expressed in a higher proportion of the other cell types *c*’relative to cell type *c*. A score close to zero, suggests that the gene is expressed in equal proportion to cell types *c*and *c*’.

##### Panel entropy score (*pes*)

This score aims to quantify the overall specificity of gene panel specificity for the cell types in a dataset by measuring spatial autocorrelation of the *gsc* scores after hierarchical clustering, weighted by the proportion of zeros. Here, a mosaic pattern arising from a low proportion of zeros and higher *gsc* scores are desirable. Such patterns indicate that the genes in a panel are predominantly cell type specific. To summarize this information, we calculate a weighted Moran’s I, which quantifies spatial autocorrelation across *gsc,c* ∈*C and g* ∈*G* for all genes and cell types. Where *N* is the number of log ratios indexed by *i* and 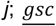 is the mean of the *gsc*; *w*_*ij*_ are the elements of the spatial weights with zeroes on the diagonal (i.e., *w*_*ij*_= 0), where we use the standardized rows of the *gsc*matrix as the weights; and *W* is the sum of all *w*_*ij*_.

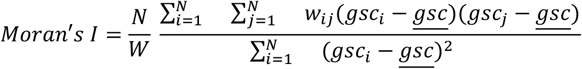

Finally, we weight Moran’s I by the proportion of zeros *p*’ to compute the panel entropy score for a given gene panel:

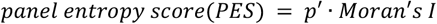

##### Variation recovery score

To quantify the ability of a gene panel to recover transcriptional variation, we calculate the Normalized Mutual Information (NMI) between the cluster assignments of the full dataset and the subsampled dataset, which is performed using *Seurat* (version 5.1.0). The purpose of this score is to evaluate how much of the transcriptional variation a panel is able to retain given it is only a subset of the total genes. A panel that retains most of the transcriptional information should be able to separate out the different cell types and states in a lower dimensional space. To quantify this, we first cluster the cells from the full dataset using all genes, referred to as Clustering *A*. Next, we cluster the dataset again using only the genes in the panel of interest, referred to as Clustering *B*. Here, the assumption is that each cluster in each clustering corresponds to a cell type or cell state. The NMI measures the similarity between the two clusterings of cells, *A* and *B*, and is calculated as

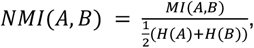

where A and B is the clustering results for the full dataset and subsampled dataset, respectively; *MI* is the mutual information between clustering *A* and clustering *B*; *H*(*A*)and *H*(*B*)are the entropy of clusterings *A* and *B*, respectively. Mutual information quantifies the amount of information shared between the two clusterings *A* and *B*, and is defined as:

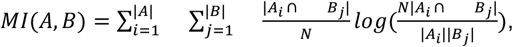

where |*A*| and |*B*| is the total number of clusters in clustering *A* and *B*, respectively; |*A*_*i*_| is the number of cells in cluster *i* of clustering *A*; |*B*_*j*_| is the number of cells in cluster *j* of clustering *B*; |*A*_*i*_∩ *B*_*j*_| is the intersection of cells between cluster *i* in *A* and cluster *j* in *B*; *N* represents the total number of cells across both clusterings.

The entropy of cluster *A* is given by:

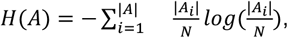

and similarly for *H*(*B)*.

#### 2.3.2. Feature diversity

##### Feature diversity score (*fds*)

To ensure an informative gene panel, it is important to maximize the diversity of information included. Thus, we aim to assess and quantify the amount of redundant information in a gene panel. We achieve this by calculating the pairwise Spearman’s correlation for all genes in the panel. For a panel *P* with *G* genes, there are *G*’ = *G*(*G* − 1)/2 pairs. We then compute the feature diversity score (*fds*), which captures the relative difference between the negative/zero correlated and positively correlated gene pairs. Gene panels with a higher number of negative/zero-correlated pairs relative to positively correlated pairs exhibit lower redundancy and higher feature diversity. Here, *p*_(_ is the Spearman’s correlation coefficient for the *i*th gene pair; we add a constant of 1 to ensure that we do not take the log of zero. The feature diversity score is defined below:

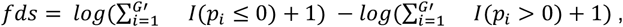

a higher *fds* score indicates greater feature diversity within a panel, whereas a lower *fds* score suggests reduced diversity and thus higher redundancy.

#### 2.3.3 Biological inference

##### Pathway diversity score (*pds*)

We summarize the pathway information contained in a gene panel by calculating various statistics, including the proportion of dissimilar pathways. Using the *enrichGO()* function from the clusterProfiler package (version 4.10.1), we perform an over-representation analysis on the genes in the panel. Significant biological pathways are identified based on an adjusted p-value threshold of less than 0.05. For each gene panel, we report the number and proportion of genes enriched in any sign significant pathway, the total number of significant pathways, the maximum q-value (false discovery rate) across all significant pathways, and a pathway diversity score (pds) to quantify the diversity in enriched pathways.

The *pds* is calculated as the one minus the proportional reduction in the number of pathways after removing pathways with a Jaccard similarity index greater than 0.7. A higher diversity score indicates that the gene panel contains a large proportion of diverse pathways, where *P* is the total number of significant pathways for the gene panel; *P*’ is the number of significant pathways remaining after removing pathways with a Jaccard similarity index <0.7; *p*_(_ is the contribution of pathway *i* to the total pathway set, and *p*_”_ is the contribution of pathway *j* to the total set of pathways remaining after removing pathways with a Jaccard similarity index greater than 0.7:

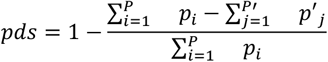

##### Cell-Cell interaction information

To evaluate the potential for cell–cell interactions within a gene panel, we first classify each gene as either a ligand or a receptor according to annotations from the CellChat database (Jin *et al*., 2021), which provides a comprehensive catalog of known ligand– receptor pairs across diverse cell types and biological systems. We then sum the number of genes in each category—ligands and receptors—to determine how enriched the panel is for each.

#### 2.3.4 Spatial information

##### Moran’s I

We capture the spatial autocorrelation of all genes within a panel by calculating Moran’s I for each gene *g*. Similar to above, let *x*_*g*)_represent the expression of gene *g*; *N* is the number of expression values indexed by *i* and 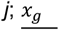 is the mean expression of gene *g* across cells; *w*_*ij*_ are the elements of the spatial weights with zeroes on the diagonal (i.e., *w*_(“_ = 0), where we use the standardized rows of the *lr* matrix as the weights; and *W* is the sum of all *w*_(“_. Moran’s I for a gene *g* is defined below:

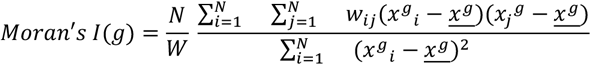

##### Nearest neighbour correlation

We calculate the correlation of the gene expression values for a panel’s genes between a cell and its nearest neighbours, using the package *scFeatures* (version 1.3.4). Briefly, scFeatures calculates the nearest neighbour correlation for each gene within a gene panel. This is calculated by taking the correlation between each cell and its nearest neighbours for a particular gene.

#### 2.3.5 Forward-compatibility

A key consideration in gene panel design is ensuring that the chosen genes are likely to remain relevant across a variety of conditions, such as different treatments, experimental groups, or perturbations. We consider this as “future versatility” or “forward compatibility”. Thus, we aim to assess this ‘future compatibility’ aspect to ensure that the panel can adapt to evolving research needs and experimental frameworks. We aim to achieve this by leveraging perturbation simulators that have recently emerged for single cell research. Here we have selected GEARS as our perturbation simulator (Roohani, Huang and Leskovec, 2024). To train the GEARS model, we utilized several public perturbation datasets (Adamson *et al*., 2016; Dixit *et al*., 2016; Norman *et al*., 2019; Replogle *et al*., 2020). Prior to training, all datasets were processed following the instructions of GEARS, such as normalizing gene expression values and filtering out low-quality cells, to ensure consistent and acceptable data across all the experiments. Once the GEARS model was trained, it was used to simulate all perturbations of each single gene existing in the training set. We focused on expression changes of genes that overlapped between the training set and the selected gene panel and these results were then collected and analysed. By focusing on the overlapping gene set, we were able to assess the sensitivity of selected genes in that panel for further validation. To summarize the perturbation potential of a gene with calculate a perturbation score, which summarizes the number and magnitude of genes it is likely to influence when perturbed.

Let *d* represent a *G*’x*G* matrix of predicted absolute expression values for *G*’ target genes which are influenced by *G* perturbed genes, calculated from training and testing on *D* datasets. The *q*-value represents the 25th percentile of the gene expression values *x* in *D*. The perturbation potential for a gene *g* is defined as:

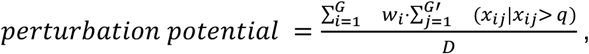

where *w*_*i*_ is a weight capturing the proportion of target genes influenced by gene *i* and have non-zero expression. This metric captures the magnitude of gene expression changes induced by target genes, normalized by the number of genes they influence. A higher perturbation potential indicates that a gene is more likely to influence the expression of other genes when perturbed. To calculate the perturbation potential of a panel, we simply take the mean of the perturbation potential scores across the genes.

#### 2.3.6 Gene importance score and overall score

##### Gene importance score

To provide a comprehensive characterization of the gene panels, A we compute an importance score for each gene within a panel, which reflects the gene’s contribution to a panel across the defined categories. Before computing the scores, we standardize each individual component from the broad categories to ensure comparability. These components are then equally weighted by default; however, users can choose to assign different weights to the components based on their specific research questions or priorities.

For a typical gene *g*, we define the gene importance score (gis) as

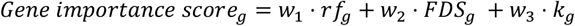

Where *rf*_*g*_ is the importance score for gene *g* output from the random forest model used for cell type classification; *fds*_*g*_ is the feature diversity score; and *k*_)_is an indicator variable, such that *k* = 1 if gene *g* is enriched in a pathway, 0 otherwise; and *w*_1_, *w*_2_, *w*_3_ are weights which can be user-specified to allow for customization based on the importance of each metric. By default, all weights are set to 0.33. Thus, gene importance scores within a panel are comparable but should not be compared across panels.

##### Weighted overall score

We calculate overall panel scores that summarize statistics across all relevant categories. These scores enable a direct comparison of characteristics between different gene panels. For a given panel *P* = {*g*_*1*_, *g*_2,_…, *g*_G_}, the weighted overall panel score is define as

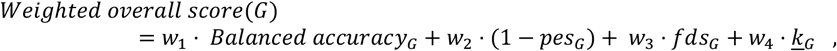

where *balanced accuracy* represents the balanced cell type classification accuracy for gene panel *G*; *pas* is the panel entropy score; *fds* is the feature diversity score; 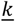 is the mean of the binary variable *k*, which indicates whether each gene in a panel is enriched in a significant pathway (*k*=1) or not (*k*=0). Since *k* is binary, 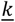 corresponds to the proportion of genes in the panel enriched in significant pathways; and *w*_*1*_, *w*_2_, *w*_*3*_, *w*_*4*_ are weights which can be user-specified to allow for customization based on the importance of each metric. By default, all weights are set to 0.25.

### 2.4 panelScope optimization algorithm

We treated the panel ingredient searching problem as a hyperparameter optimization problem. From this perspective, each position in the designed gene panel would be a hyperparameter in which all genes are candidates and thus, the searched panel would be a combination of genes. From an initial randomly picked state, the optimization goal is to find gene replacements gradually, continuously and efficiently with better desired characteristics.

#### 2.4.1 Objective function module

The modular architecture of the panelScope optimization algorithm renders it great flexibility and versatility. panelScope could be divided into two main parts, i.e., the objective function module and the tuning module. We used the normalized even-weighted sum of all metrics described above in Section 2.3 as our objective function. Its value ranges from 0 to 1, where 0 represents a not-qualified panel and 1 represents a perfect panel. As to hyperparameter space exploration, we chose approaches employing multi-trial strategy to do the optimization. Although conventional, the multi-trial strategy is sharply straightforward and effective. This strategy independently constructs and evaluates each gene panel sampled from the predefined gene set, which allows researchers to assess the characteristics of gene panels based on their own merit and give researchers the ability to gain comprehensive insights, without any interference or shared information from other panels.

#### 2.4.2 Tuning module

Due to its modular architecture, every component of panelScope could be substituted. As mentioned before, we chose the multi-trial strategy for hyperparameter space exploration. We tried several different Bayesian and heuristic tuning algorithms suitable for this optimization such as Tree-structured Parzen Estimator (TPE, the representative of Bayesian tuners) (Feitosa Neto and Canuto, 2021), Anneal algorithm (the representatives of heuristic tuners) and Evolution algorithm. Our experience shows Evolution has the greatest potential. Evolution algorithm operates by initially generating a population of gene panels that are randomly sampled from the predefined search space, i.e., the gene set. During each generation of the evolutionary process, the panels within the population are evaluated based on the weighted score of their characteristics, with the better-scoring panels being selected for further refinement. The mutation, i.e., the process of refinement, involves making slight modifications to the selected panels, such as replacing, adding or removing genes. The mutated panels are then used to form the subsequent generation. This iterative and adaptive panel modification would result in great characteristics improvement in the end.

## 3. Results

### 3.1 Multi-view gene panel characterization presents a comprehensive view for a pre-defined gene panel

We present panelScope, a modular framework for multi-view gene panel characterization, accessible via a user-friendly web application (Figure 1A). panelScope characterizes gene panels using metrics grouped into five broad categories, with flexibility for future extensions (Figure 1B). To enhance usability, panelScope generates overall scores at both the gene and panel levels, summarizing performance across these categories. This enables users to identify and rank the genes that contribute the most valuable information within a panel, while the panel level scores can be used to compare across the different panels. By providing a detailed overview of each panel’s strengths and weaknesses, panelScope allows researchers to make informed decisions when designing or refining their panels.

**Figure 1.**
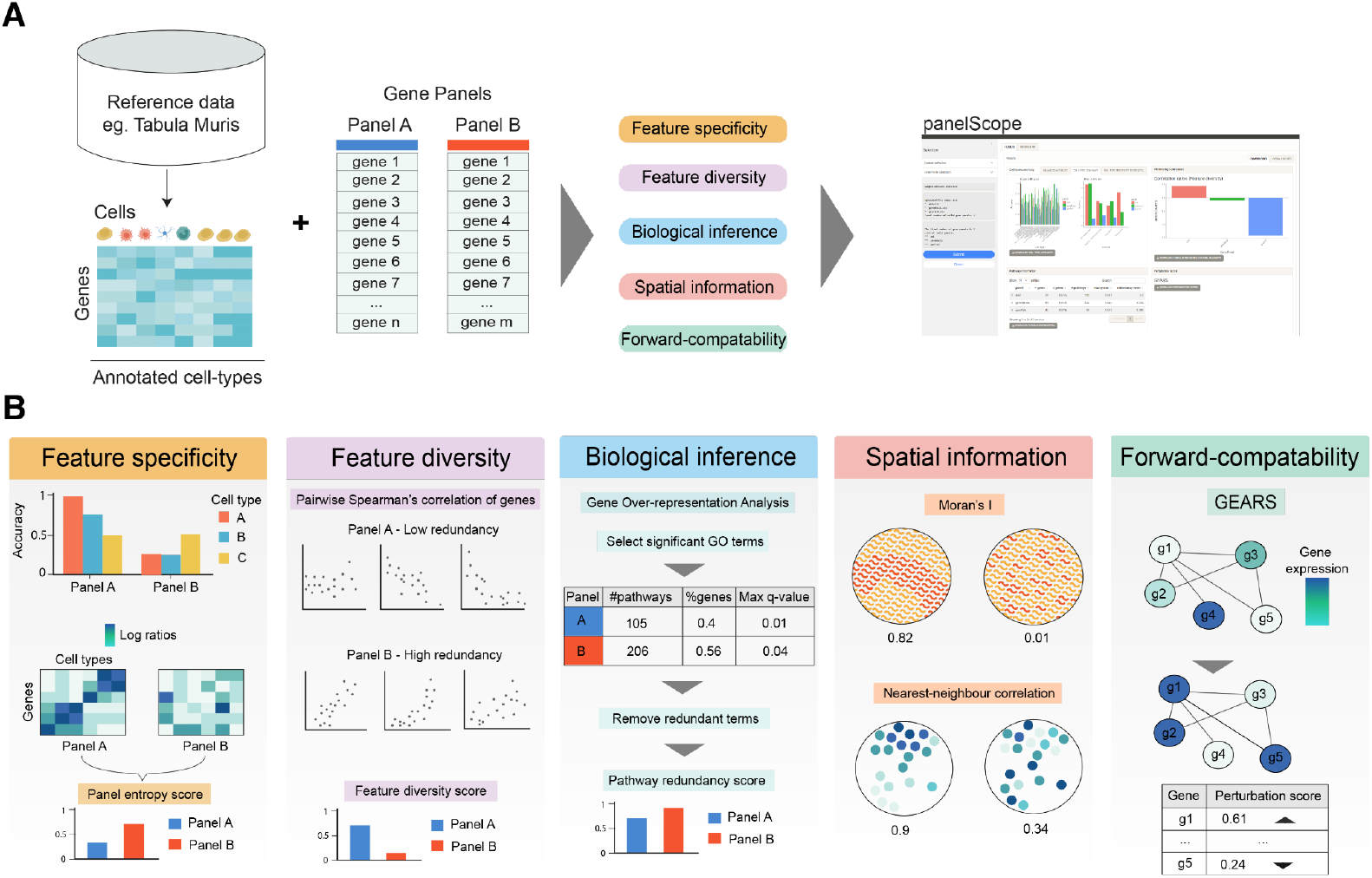
A) Workflow of gene panel characterization using panelScope. B) Description of the five categories of metrics used to characterize a given gene panel: Feature specificity, feature diversity, knowledgebase, spatial information, and prediction.

Additionally, we have developed a genetic algorithm that utilizes these metrics to design and optimize gene panels tailored to specific research objectives. To further support customization, panelScope allows users to assign adjustable weights for the four categories, allowing users to adjust the scoring system based on their specific research questions. This feature ensures that panel evaluations are tailored to the specific needs of users, making the framework relevant for a range of applications. Whether the focus is on maximizing pathway coverage, identifying cell type-specific markers, or ensuring broad feature diversity, users can fine-tune the optimization algorithm to align with their interests.

### 3.2 Application of panelScope to AML to illustrate cell type-specificity of panels

As a case study we applied our framework to design a panel guided by a dataset specific to acute myeloid leukemia (Beneyto-Calabuig *et al*., 2023). We assessed the performance of each panel across the categories, starting with their ability to classify cell types in the reference dataset using genes from the respective panels. Classification performance was quantified using balanced accuracy and accuracy (Figures 2A,-C).

**Figure 2.**
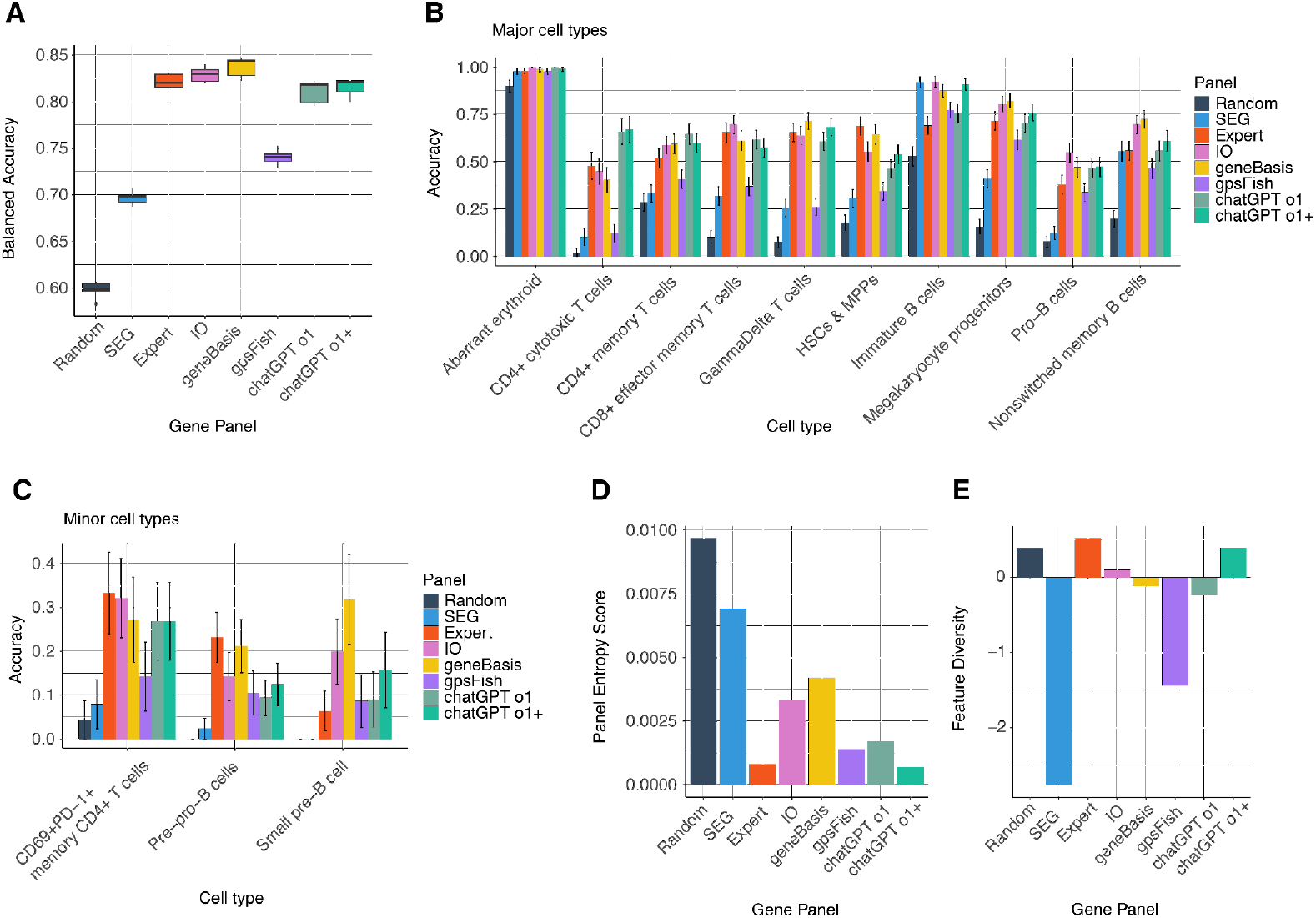
A) Boxplots of balanced accuracy. This represents the overall ability of a gene panel to capture both major and minor cell types in the tested dataset. B) Barplot of a gene panel’s ability to capture major cell types in the tested dataset. Error bars represent one standard error above and below the mean accuracy. C) Similar to panel B) but stratified for minor cell types. D) Panel entropy redundancy score for each gene panel. A lower score indicates a higher proportion of cell type specific genes in panel. Thus, the lower the more desirable. E) Bar plot of feature diversity for each gene panel. A positive score indicates higher diversity of features while a negative score indicates lower diversity of features (high redundancy).

In terms of overall performance in capturing cell types, the panels curated by our in-house expert, *IO, geneBasis*, and *chatGPT* models achieved similar balanced accuracy. Conversely, the *Random* and *SEG* panels performed relatively worse, as expected, due to their lower specificity. Interestly, the *Random* and *SEG* panels still maintained a balanced accuracy greater than 0.5, suggesting that some genes in these panels are cell type specific. This was surprising as stably expressed genes are typically genes that maintain consistent expression levels across cell types and are related to basic cellular functions, such as glycolysis (Lin *et al*., 2019). Nevertheless, under some conditions, stably expressed genes can be perturbed in a cell type specific manner. For example, a small number of genes in the *SEG* panel, such as BIRC5, have been implicated in cancer cell survival across multiple cancers (Xu *et al*., 2021; Li *et al*., 2024), suggesting that these stably expressed genes are perturbed in our particular case study, resulting in some cell type specificity.

To provide a more detailed view of a panel’s cell type specificity, we assess whether a panel contains the appropriate genes to accurately capture both major and minor cell populations as shown in Figures 2B and 2C. For all panels, we observed high accuracy in capturing specific major cell types, such as the aberrant erythroid cells, with an accuracy greater than 0.8 (Figure 2B). However, the *Random* and *SEG* panels demonstrated poor performance in capturing minor cell types, emphasizing the increased importance of careful panel design when targeting rare or less abundant populations.

To assess feature specificity, we developed a panel entropy score, which reflects the relative proportion of non-cell type-specific genes within a panel (Figure 2D). First, we compute a gene specificity score (gsc) for each gene, defined by the log ratio of the proportion of cells expressing that gene in a target cell type relative to all other cell types. Genes with gsc values above zero are considered cell type-specific, while those close to zero indicate minimal or no specificity. We then perform hierarchical clustering on the gsc and visualize them as a heatmap, where spatial patterning reflects enrichment of cell type-specific genes (low entropy), and a lack of patterning indicates the presence of many non-cell type-specific genes (high entropy) (Figure 3A). We quantify these patterns using a weighted Moran’s I of the gsc, where higher autocorrelation reflects a higher number of cell type-specific genes in the panel. This measure is further adjusted by the proportion of genes with gsc scores of zero, ensuring a comprehensive assessment. Therefore, a lower panel entropy score indicates a panel enriched with cell type-specific genes and as expected, the *Random* and *SEG* panels exhibited minimal cell type specificity (Figure 2D). Additionally, we measure how much transcriptional variation each panel recovers using Normalized Mutual Information (NMI) (Figure 3B). We find that the *geneBasis* panel captures more transcriptional variation than the other panels, likely because it explicitly optimizes for preserving the global transcriptional diversity of the reference data.

**Figure 3.**
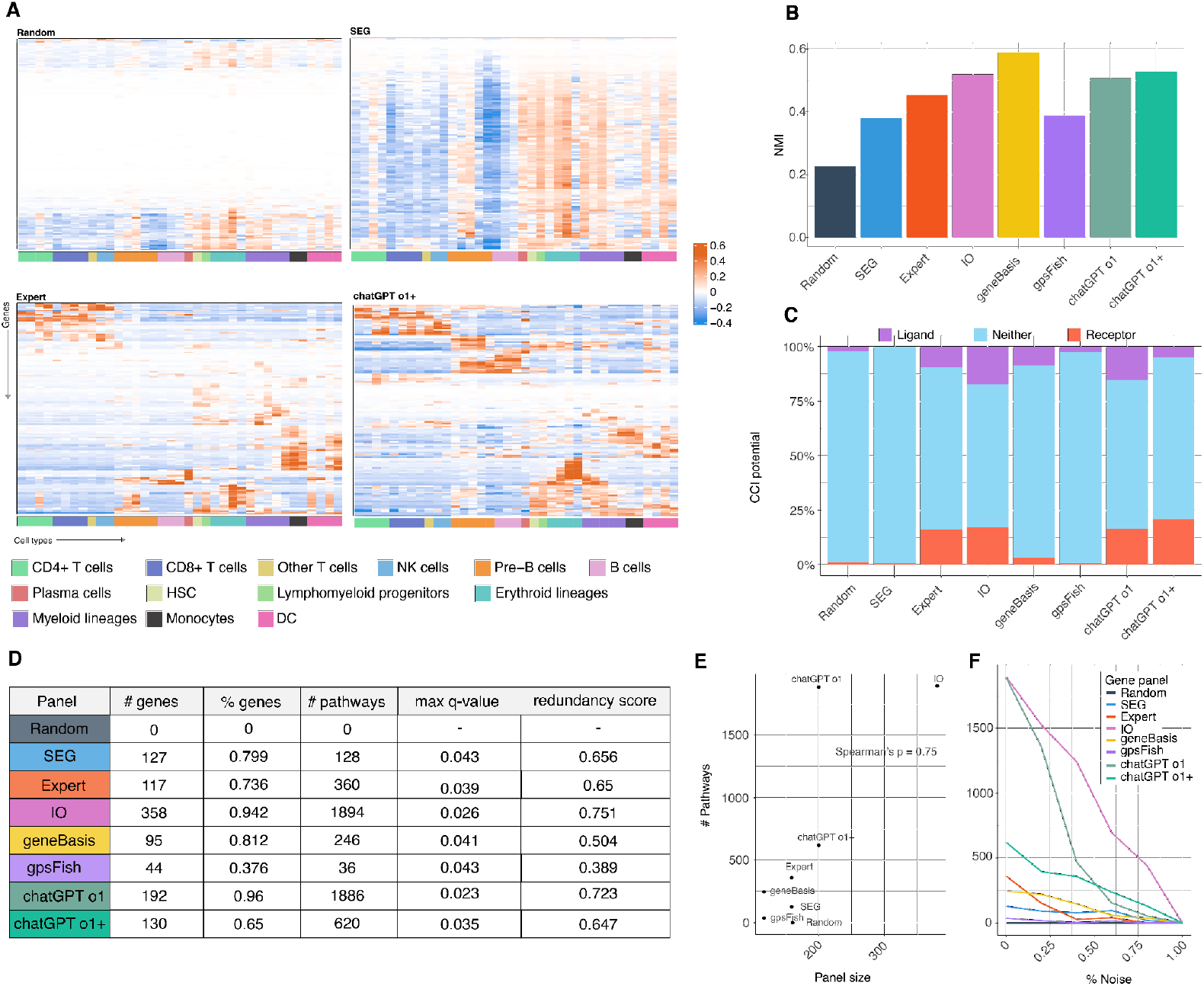
**A)** Heatmaps of gene specificity scores. Rows represent genes while columns represent cell types, which have been grouped into their major classes. Top row of heatmaps display undesirable panels as they show non-specificity (*Random*) or high specificity for a select few cell types (*SEG*), while the bottom row show desirable panels (*Expert-curated, chatGPT o+*). **B)** NMI represents the amount of transcriptional variation that panels are able to retain **C)** Stacked barplot showing the number of ligands, receptors, and genes that belong to neither category for each panel **D)** Table of pathway information for each gene panel, all metrics are calculated on the significant pathways (adjusted p-value < 0.05). # genes: number of genes from panel that were enriched in significant pathways; % genes: proportion of genes in panel that were enriched in significant pathways; # pathways: number of significant pathways for a given gene panel; max q-value: maximum q-value (false positive rate) out of all significant pathways; redundancy score: relative change in number of significant pathways when removing pathways with a Jaccard similarity index > 0.7. **E)** Scatter plot to investigate the correlation between the number of genes in a gene panel and the number of significant pathways and the proportion of genes enriched in significant pathways. **F)** Scatter plot of the number of significant pathways for each panel as varying proportions of random genes (noise) is added.

### 3.3 Feature Diversity Score: Panels with orthogonal information (e.g., *Random* and *Expert-curated* panels) exhibited higher diversity compared to *SEG*, which contained highly correlated genes

Here, we sought to summarize the redundancy or diversity of features within a panel. While the panel entropy score measures the cell type specificity of a panel, the feature diversity score evaluates the extent to which a panel provides orthogonal (non-redundant) information relative to redundant information (Figure 2E). This score is calculated as the number of gene pairs with a Spearman’s correlation of zero or below relative to those with a positive correlation. A higher feature diversity score, therefore, reflects greater diversity of information provided by the panel’s features (Figure 2E). The SEG panel exhibited the highest number of positively correlated genes, indicating significant co-expression among housekeeping genes. In contrast, the *Random, Expert-curated*, and *chatGPT o1+* panels demonstrated the highest feature diversity. While this may seem counterintuitive for the *Random* set of genes, the selection of random genes is unlikely to result in a high proportion of positively correlated pairs. Instead, many gene pairs in the *Random* panel are likely to have zero or negative correlations, contributing to its high diversity score.

### 3.4. Biological inference

The Biological inference category aims to evaluate how much meaningful biological information each panel is likely to provide. To achieve this, we visualize the number of ligands and receptors in each panel (Figure 3C) and assess the pathway information they capture through gene over-representation analysis (Figure 3D). As expected, the *Random* and *SEG* panels contained the fewest ligands and receptors. Interestingly, we observed that the *Expert-curated, chatGPT o1*, and *chatGPT o1+* panels had the highest proportion of ligands and receptors (26%, 31%, and 26%, respectively), outperforming algorithm-based methods such as *geneBasis* (11%) and *gpsFISH* (0.3%). The higher ligand and receptor representation in the *Expert-curated* and *IO* panels can be attributed to leveraging a hybrid approach to panel design (computational and manual). Surprisingly, despite lacking any manual curation, the *chatGPT* panels perform comparably - and sometimes exceed - the hybrid-designed panels (Figure 3C). This advantage may arize from the reasoning capabilities of large language models and their exposure to vast amounts of pre training data, which seem to enhance ligand and receptor selection.

To evaluate biological pathway information within a panel, we use several metrics: the number and proportion of genes enriched in significant pathways, the total number of significant pathways, the maximum q-value among significant pathways, and a pathway diversity score (Figure 2D). The pathway diversity score reflects the degree of non-overlapping pathways, with higher scores indicating higher pathway diversity due to a lower proportion of shared genes. We noticed that some of these metrics appeared to be influenced by panel size (Figure 3E). For example, the number of significant pathways enriched in a panel is positively correlated with panel size (Spearman’s correlation=0.75) (Figure 3E). However, we found that this relationship depends on the inclusion of subsets of genes that share biologically meaningful relationships. For instance, the *Random* panel, which lacks this property, showed no pathway enrichment. To further investigate, we calculated these metrics after introducing varying proportions of random genes (noise) into each panel. As the proportion of random genes increased, the number of significant pathways decreased across all panels (Figure 3F). This demonstrates that the positive correlation between panel size and the number of significant pathways is confounded by the proportion of genes that share biological relationships (Figure 3F).

### 3.5 Forward compatibility and spatial information

We offer two optional categories: forward compatibility and spatial information. These categories are optional due to potential users’ computational limitations and current data availability but can be utilized when sufficient data (e.g. spatial data) is available and may be extended in the future.

Forward Compatibility. The aim of the metrics in the forward compatibility category are to identify panels that contain genes which have the highest potential for perturbing other genes given they are perturbed themselves. Recently, various methods have been developed to predict gene perturbations from single-cell datasets via simulations. We leveraged one such approach-GEARS. We applied GEARS across five datasets, which included multiple perturbation conditions, to simulate and estimate the perturbation potential of individual genes before averaging them to compute the perturbation potential for each of the eight panels tested(Figure 4A). Panels with higher scores are predicted to contain genes that, when perturbed, are likely to impact a greater number of downstream genes across various conditions. Of all the tested panels, the *SEG* panel was predicted to contain genes with the highest perturbation potential. This is expected as stably expressed genes are typically involved in essential and fundamental cellular processes and (Eisenberg and Levanon, 2013). Thus, perturbing such genes can lead to significant cellular effects, including cellular death and widespread transcriptional changes (Blomen *et al*., 2015).

**Figure 4.**
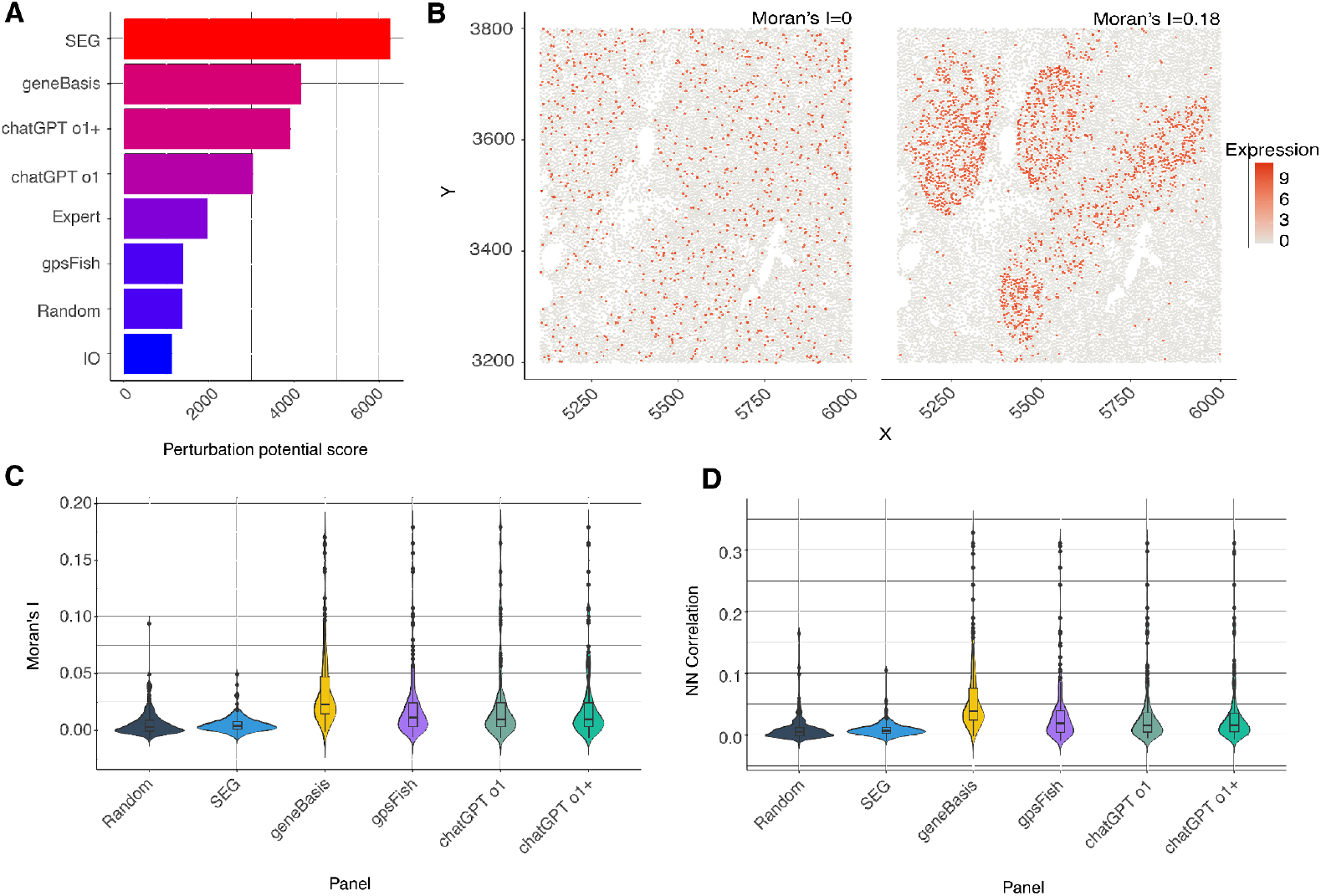
**A)** Panels are ranked in descending order based on their perturbation potential score. Higher scoring panels contain genes that are more likely to be perturbed themselves **B)** Representative spatial plots of two genes with contrasting spatial autocorrelation. The left plot shows a gene (PROCR) with random expression across the tissue section (low Moran’s I), while the right plot shows a gene (CXCL13) with a pronounced clustered pattern of expression (high Moran’s I). Red dots represent higher expression levels, and white/gray areas indicate lower or no expression. **C)** Violin plots of Moran’s I across all tested gene panels. Each violin plot visualizes the distribution, median, and spread of the respective spatial statistic across all genes in the panel. **D)** Violin plots of nearest‐neighbor (NN) correlation for each gene panel, providing an alternative metric of spatial clustering.

Spatial Information. Another important characteristic of a panel, particularly for spatial transcriptomics, is whether a panel is enriched for genes that exhibit spatial variation. Genes lacking spatial variation in a given tissue context may not provide additional spatial insights beyond single-cell RNA-seq and thus might not be as valuable to include in the panel (Figure 4B). To evaluate spatial information, we calculate Moran’s I and Nearest Neighbour Correlation for each gene in a panel (Figures 4C,D). As expected we observed overall lower Moran’s I and Nearest Neighbour Correlation for the *Random* and *SEG* panels, indicating the determination of these panel didn’t account for any spatial information. Conversely, the *geneBasis* panel showed higher Moran’s I and Nearest Neighbour Correlation compared to all other characterized panels, presumably due to its algorithm’s focus on preserving transcriptional variation, which may be associated with capturing cell type markers, which can have spatial variability due to the fact that cell types are distributed non-uniformly across complex tissues (Figure 4C,D) (Maynard *et al*., 2021; Shang, Wu and Zhou, 2025). Furthermore, this metric is a gene based meteric and allowing users to manually refine the panel based on characteristics of individual genes.

### 3.6 panelScope leverages the multi-view panel characteristics to create gene panel panelScope

We illustrate the panelScope gene panel selection method in Figure 5A. As we can see, the algorithm conducts an iterative optimization process which mainly contains four steps. Initialization is the very beginning step where the algorithm will randomly select a number of gene panels (default as 50) from the input dataset to form the initial population. Then, the algorithm will conduct an evaluation-selection-generation round for a number of times (default as 5000). In each round, evaluation scores quantify how well candidate gene panels perform on the selected characteristic metrics, where higher scores indicate better‐performing panels. Then the algorithm would randomly pick better gene panels, use them to generate new panels and replace those that do not perform well. Once the evaluation threshold was reached or the optimization process had been conducted for certain rounds, our algorithm would stop and choose the best panel of the history as the final results. By these, our genetic algorithm rapidly converges on a high‐performing solution, then continues to refine it incrementally. The consistent maximum‐score curves across rounds reflect the robustness of this optimization, suggesting that panelScope effectively balances exploration and exploitation.

**Figure 5.**
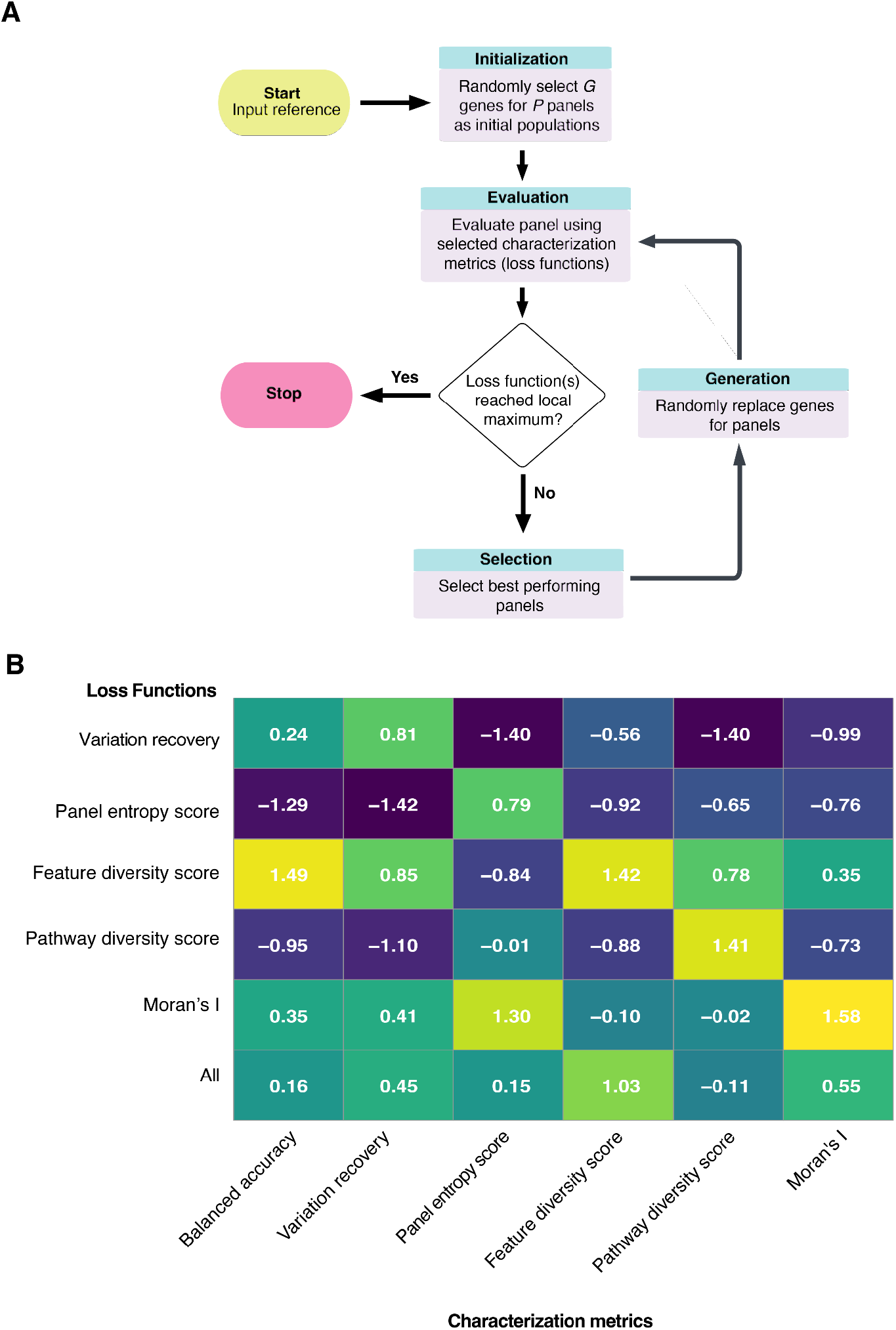
**A)** Algorithm illustration of gene panel selection. An iterative optimization process containing 4 steps is evolved. At the initialization step, a number of gene panels (default as 50) are randomly selected from the input dataset as initial population. Then, an evaluation-selection-generation loop would be repeated for a number of times (default as 5000). We use scoring functions related to feature diversity, pathway diversity, panel entropy, spatial specificity and variation recovery as our evaluation metrics. The algorithm would randomly pick better performed gene panels and use them to generate new panels to replace the old, not-well-performed ones. **B)** Heatmap of standardized scores for six curated gene panels, each optimized under a different *loss function*, evaluated across multiple *characterization metrics*. Rows correspond to the chosen loss function (e.g., Variation recovery, Feature diversity score), while columns show the various metrics used for panel characterization (e.g., Balanced accuracy, Pathway diversity score). Color‐coded cells represent the performance of each panel on the respective metric, with higher or lower values (depending on the metric) indicated by more saturated colors. Panel entropy score was multiplied by −1 for interpretability.

We grouped our characterization metrics into five broad categories—feature specificity, feature diversity, biological inference, spatial information, and forward compatibility—and monitored optimization progress under each metric type both independently and in combination (Supplementary 3-4). This approach revealed how different optimization strategies evolved over time, and we compared those results with the final optimized panel shown in Figure 5B. We found that the variation recovery and panel entropy score loss functions provided genes with information unique to their respective categories, while leveraging feature diversity consistently produced stronger optimization gains. Thus, the panel selected using the feature diversity loss function appeared to provide positive information across multiple categories (Figure 5B).

## 4. Discussion

Here, we have presented panelScope, a comprehensive gene panel characterization framework that can be used to characterize a gene panel from multiple perspectives including cell type specificity, feature diversity, biological inference, spatial information, and forward-compatibility. We also provide two accompanying tools to enhance users’ engagement with this framework and illustrate how these tools can support gene panel design decisions. To achieve this, we developed an interactive web tool that enables scientists to understand the characteristics of a given gene panel and we illustrate how this can be used to design more informative panels. In parallel, we developed an optimization algorithm that incorporates a differential weighting scheme or emphasis on panel characteristics, allowing for custom panel design.

Currently, there is no framework to quantitatively assess the informativeness of a gene panel, making it difficult to make informed decisions when designing or customizing a gene panel. panelScope enables researchers to comprehensively characterize a gene panel and determine whether the chosen genes provide information relevant to their research objectives. panelScope not only offers performance measures across five categories, but the framework also computes gene-level scores, reflecting the level of information each gene contributes across these categories. This allows researchers to fine-tune their gene panel and make informed decisions about which genes to include or exclude from their panels. Importantly panelScope, is not limited to characterizing manually selected panels and can also assess gene panels derived from any panel selection method, including commercially available panels.

While the effort in the community to date has focused on determining the optimal gene panel, it is important to recognize that the optimality of a gene panel is criteria-dependent. The optimality of a gene panel depends on the specific research objectives it is designed to address. For instance, a panel could select genes that are key markers for immune cell types in a particular cancer, such as CD8+ T cells or tumor-associated macrophages (TAMs) in lung cancer, to assess immune infiltration and cancer immunotherapy targets (Chen and Mellman, 2017). Alternatively, a panel could focus on signaling pathways like PI3K-AKT or RAS-RAF-MEK, which are relevant across multiple cancers, where capturing diverse pathways is desirable (Roberts and Der, 2007; Engelman, 2009). Naturally, given the limited number of genes that can be selected, there is a trade-off between these objectives. Thus, our interactive web tool enables users to explore these trade-offs, relating this information to the prioritization of specific objectives over others, allowing them to make quantitatively informed decisions on panel design. This flexibility ensures that researchers can tailor their gene panels to meet their specific research goals, whether they are focused on addressing multiple research questions or homing in on specific areas of interest.

An important factor in designing a gene panel is ensuring that the selected genes are likely to be forward compatible, meaning they will be differentially expressed across various treatments, experimental groups, or perturbation conditions. This aspect of “future compatibility” ensures the panel remains relevant over time, allowing it to adapt to evolving research needs and experimental designs. Our current strategy leverages simulations with the GEARS framework, though this component can be replaced with improved simulation strategies in the future. Emerging methods for perturbation prediction utilize existing datasets from drug and genetic perturbations to forecast cellular responses to unseen changes. These approaches incorporate recent advances in AI and multi-omics data, such as scGen, which uses autoencoders for latent space shifts; CellOT, which applies optimal transport theory; CellOracle, which integrates chromatin accessibility for genetic perturbations; and PerturbNet, which uses both genetic and chemical structure data, (Lotfollahi, Wolf and Theis, 2019; Yu and Welch, 2022; Bunne *et al*., 2023; Kamimoto *et al*., 2023). As these algorithms improve in the coming years, the metrics associated with this category in panelScope will also evolve, providing increasingly accurate and refined predictions for future compatibility.

Although panelScope was designed to characterize and evaluate gene panels designed for targeted spatial transcriptomics, many of the characterization categories are not dependent on spatial statistics, thus making panelScope generalisable to any technology requiring targeted gene panel selection, such as nCounter or Ion AmpliSeq. This may seem counterintuitive, however spatial metrics have limited applicability for characterizing gene panels due to the scarcity of comprehensive spatial datasets that can be leveraged for this purpose. Instead, we have intentionally designed multiple spatially agnostic metrics that can achieve comparable goals, while taking advantage of the vast availability of single-cell transcriptomic data. However, overtime, this will change as more spatial datasets become available. Importantly, panelScope can be extended to incorporate multi-omics data, such as proteomics, as these other molecular platforms often offer complementary information that will enhance the comprehensiveness of the panel selection. This would allow the framework to capture a broader and more comprehensive view of biological characteristics.

Going forward, the accompanying genetic algorithm for gene panel design can be further improved through the incorporation of reinforcement learning techniques. Although much more computational resources would be needed, reinforcement learning would enable the framework to learn from feedback throughout the selection process, continuously refining gene panels based on the selected evaluation metrics. This approach would have a chance to enable a more adaptive and personalized optimization process, where the system would evolve and improve with each iteration based on real-time feedback.

## 5. Conclusion

In conclusion, we have developed a comprehensive framework for gene panel characterization across multiple views. This is crucial for allowing transparent comparisons between gene panels and enables researchers to make informed decisions when designing gene panels. In addition, we have used these metrics to develop a genetic algorithm for panel selection. Unlike other panel selection methods, the panelScope optimization algorithm efficiently balances exploration and exploitation, continually refining gene panel selection while avoiding premature convergence.

## Data Availability

The panelScope web application can be accessed at https://shiny.maths.usyd.edu.au/Marni/Test/panel_selection/. The panelScope optimization algorithm can be accessed at https://github.com/SydneyBioX/panelScope/tree/main.

## Acknowledgments

The authors thank all their colleagues, particularly at The University of Sydney, members from the Sydney Precision Data Science Centre for their support and intellectual engagement. Special thanks to Ellis Patrick and Lijia Yu for their contribution in our weekly discussions.

## Author Contribution

J.Y. conceived, designed and funded the study. DK led the metric development and evaluation with input from A.N and guidance from J.Y and P.Y; W.D led the optimization algorithm and analysis with input from D.K and A.N. The implementation of the package was done by W.D and the construction of the interactive web application was done by M.T. DK led the manuscript writing and all authors edited, reviewed, and approved the manuscript.

