## Supplementary material for "Multi-view gene panel characterization for spatially resolved omics"

### Supplementary 1

Prompt 1

I am designing a gene panel consisting of 200 genes in the context of Acute Myeloid Leukemia (AML). The goal is to optimize the panel based on the following criteria:

- Cell type specific markers to capture major and minor cell types in AML.
- Transcriptional variation, try to capture as much of the transcriptional variation possible.
- Feature diversity, ensuring minimal redundancy among positively correlated features.
- Inclusion of diverse pathways to capture broad biological relevance.
- Representation of ligands and receptors, crucial for cell-cell communication.
- Inclusion of spatially variable genes, capturing spatial heterogeneity in AML.

Please design a gene panel of 200 genes based on these criteria. Provide the gene list in one continuous line, with each gene enclosed in quotation marks and separated by commas.

Response 1

“CD34”,”KIT”,”FLT3”,”GATA2”,”RUNX1”,”MECOM”,”MEIS1”,”HOXA9”,”BMI1”,”EZH2”,
“MPO”,”CEBPA”,”CEBPE”,”SPI1”,”CSF1R”,”CSF3R”,”ITGAM”,”LYZ”,”S100A8”,”S100A9”,
“CD14”,”FCGR3A”,”ITGAX”,”CD68”,”MRC1”,
“GYPA”,”HBB”,”HBA1”,”ALAS2”,”KLF1”,”EPOR”,”SLC4A1”,”FECH”,
“ITGA2B”,”GP1BA”,”PF4”,”NFE2”,”TUBB1”,
“MS4A1”,”CD19”,”CD79A”,”CD79B”,”PAX5”,”EBF1”,”IGHM”,”IGHD”,”CD22”,”VPREB1”,
“CD3D”,”CD3E”,”CD3G”,”CD4”,”CD8A”,”CD8B”,”CD2”,”CD5”,”CD7”,”CD28”,”CTLA4”,”PDCD1”,”ICOS”,”TRAC”,”TRBC1”,
“NCR1”,”KLRD1”,”KLRC1”,”KLRC2”,”KIR2DL3”,”KIR3DL1”,”GNLY”,”NKG7”,”PRF1”,”GZMB”,
“CD1C”,”CLEC9A”,”XCR1”,”IRF8”,”BATF3”,”FCER1A”,”CLEC10A”,”SIRPA”,
“SDC1”,”IRF4”,”PRDM1”,”XBP1”,
“LILRA4”,”IL3RA”,”TCF4”,”SPIB”,”IRF7”,
“GATA1”,”IL5RA”,”CPA3”,”PRG2”,”KITLG”,
“NPM1”,”DNMT3A”,”TET2”,”IDH1”,”IDH2”,”KRAS”,”NRAS”,”TP53”,”CBFB”,”WT1”,”KMT2A”,”MLLT3”,”PML”,”RARA”,”SF3B1”,
“CD40”,”CD40LG”,”CXCL12”,”CXCR4”,”PDCD1LG2”,”CD274”,”CD80”,”CD86”,”ICAM1”,”VCAM1”,”SELE”,”SELP”,”SIRPB1”,”MICA”,”MICB”,”NOTCH1”,”JAG1”,”DLL1”,”TGFB1”,”TGFBR1”,”TGFBR2”,
“WNT3A”,”WNT5A”,”FZD1”,”CTNNB1”,”AXIN1”,
“JAK2”,”STAT3”,”STAT5B”,”SOCS1”,”SOCS3”,
“CD44”,”VIM”,”FN1”,”MMP9”,”ANGPT1”,”ANGPT2”,”COL1A1”,”COL3A1”,”LAMC2”,”ITGB1”,
“MYB”,”GFI1”,”BCL11A”,”FOXO3”,”CEBPD”,”IRF1”,”ETS1”,”IKZF1”,”NFIA”,”RUNX2”,
“BCL2”,”MCL1”,”BAX”,”BAK1”,”CASP3”,”CASP8”,”CDKN1A”,”CDKN2A”,”MDM2”,”RB1”,

“GAPDH”,”LDHA”,”PFKP”,”HK1”,”G6PD”,”IDH3A”,”SDHA”,”NDUFA9”,”COX4I1”,”ATP5F1B”,
“CXCL8”,”CCL2”,”CCL3”,”CCL5”,”IL1B”,”IL6”,”TNF”,”CD70”,”TNFRSF1A”,”IL10”,
“ASXL1”,”BCOR”,”PHF6”,”ARID1A”,”SMARCA4”,”CREBBP”,”EP300”,”CBX5”,”SUZ12”

Prompt 2

I am designing a gene panel consisting of 200 genes in the context of Acute Myeloid Leukemia (AML). The goal is to optimize the panel based on the following criteria:

- Cell type specific markers, where the cell types of interest are: Aberrant erythroid, CD4+ cytotoxic T cells, CD4+ memory T cells, CD4+ naive T cells, CD56brightCD16- NK cells, CD56dimCD16+ NK cells, CD69+PD-1+ memory CD4+ T cells, CD8+ central memory T cells, CD8+ effector memory T cells, CD8+ naive T cells, CD8+CD103+ tissue resident memory T cells, Class switched memory B cells, Classical Monocytes, Conventional dendritic cell 1, Conventional dendritic cell 2, Early erythroid progenitor, Early promyelocytes, Eosinophil-basophil-mast cell progenitors, Erythro-myeloid progenitors, GammaDelta T cells, HSCs & MPPs, Immature B cells, Late erythroid progenitor, Late promyelocytes, Lymphomyeloid prog, Mature naive B cells, Megakaryocyte progenitors, Myelocytes, Non-classical monocytes, Nonswitched memory B cells, Plasma cells, Plasmacytoid dendritic cell progenitors, Plasmacytoid dendritic cells, Pre-B cells, Pre-pro-B cells, Pro-B cells, Small pre-B cell
- Transcriptional variation, try to capture as much of the transcriptional variation possible.
- Feature diversity, ensuring minimal redundancy among positively correlated features.
- Inclusion of diverse pathways to capture broad biological relevance.
- Representation of ligands and receptors, crucial for cell-cell communication.
- Inclusion of spatially variable genes, capturing spatial heterogeneity in AML.

Please design a gene panel of 200 genes based on these criteria. Provide the gene list in one continuous line, with each gene enclosed in quotation marks and separated by commas.

**Response 2**

“GYPA”,”HBB”,”HBA1”,”KLF1”,”EPOR”,”CD4”,”GZMB”,”PRF1”,”FASLG”,”EOMES”,”CD4”,”CD44”,”IL7R”,”CD28”,”CXCR5”,”CD4”,”TCF7”,”CCR7”,”LEF1”,”SELL”,”NCAM1”,”NKG7”,”IL2RB”,”ZEB2”,”KIT”,”NCAM1”,”FCGR3A”,”PRF1”,”GNLY”,”KIR2DL3”,”CD4”,”CD69”,”PDCD1”,”IL2RA”,”CXCR3”,”CD8A”,”TCF7”,”IL7R”,”SELL”,”CD27”,”CD8A”,”GZMA”,”GZMB”,”PRF1”,”GNLY”,”CD8A”,”CCR7”,”LEF1”,”TCF7”,”PTPRC”,”CD8A”,”ITGAE”,”CD69”,”ITGA1”,”CXCR6”,”CD19”,”MS4A1”,”AICDA”,”CD27”,”IGHG1”,”CD14”,”LYZ”,”S100A8”,”S100A9”,”FCGR1A”,”CLEC9A”,”XCR1”,”BATF3”,”IRF8”,”CADM1”,”CD1C”,”FCER1A”,”CLEC10A”,”SIRPA”,”IRF4”,”GATA1”,”TAL1”,”KLF1”,”AHSP”,”ALAS2”,”MPO”,”CD33”,”CEBPA”,”GFI1”,”LYZ”,”GATA2”,”PRG2”,”CPA3”,”IL5RA”,”ENPP3”,”CD34”,”GATA2”,”GATA1”,”CSF1R”,”SPI1”,”TRDV1”,”TRDV2”,”TRGC1”,”TRGC2”,”CD3D”,”CD34”,”KIT”,”FLT3”,”MEIS1”,”HOXA9”,”CD19”,”IGM”,”IGHD”,”VPREB1”,”PAX5”,”GYPA”,”FECH”,”SLC4A1”,”TMOD1”,”GATA1”,”MPO”,”CEBPE”,”PRTN3”,”ELANE”,”CD33”,”MEF2C”,”DNTT”,”RAG1”,”RAG2”,”IL7R”,”CD19”,”MS4A1”,”IGHD”,”CR2”,”BANK1”,”ITGA2B”,”GP1BA”,”PF4”,”NFE2”,”TUB

B1”,”MPO”,”ELANE”,”CEBPE”,”PRTN3”,”MMP8”,”FCGR3A”,”NR4A1”,”CX3CR1”,”CD14”,”ITGAL”,”CD19”,”MS4A1”,”CD27”,”FCER2”,”IGHM”,”SLAMF7”,”CD38”,”XBP1”,”PRDM1”,”IRF4”,”TCF4”,”IL3RA”,”LILRA4”,”SPIB”,”IRF8”,”IL3RA”,”LILRA4”,”TCF4”,”IRF7”,”GZMB”,”VPREB1”,”IGLC2”,”IGLL1”,”CD79A”,”EBF1”,”KIT”,”FLT3”,”EBF1”,”TCF3”,”IGLL1”,”PAX5”,”EBF1”,”CD79A”,”CD79B”,”VPREB3”,”VPREB3”,”CD24”,”PAX5”,”RAG1”,”RAG2”,”NPM1”,”IDH1”,”IDH2”,”WT1”,”TP53”,”DNMT3A”,”TET2”,”CBFB”,”KMT2A”,”RARA”,”CXCL12”,”CXCR4”,”CD274”,”PDCD1LG2”,”HLA-DRB1”

### Supplementary 2

<u>Prompt 1</u>

I am designing a gene panel consisting of 200 genes in the context of lymph node cancer in humans. The goal is to optimize the panel based on the following criteria:

- Cell type specific markers to capture major and minor cell types in lymph node cancer.
- Transcriptional variation, try to capture as much of the transcriptional variation possible.
- Feature diversity, ensuring minimal redundancy among positively correlated features.
- Inclusion of diverse pathways to capture broad biological relevance.
- Representation of ligands and receptors, crucial for cell-cell communication.
- Inclusion of spatially variable genes, capturing spatial heterogeneity in the lymph node.

Please design a gene panel of 200 genes based on these criteria. Provide the gene list in one continuous line, with each gene enclosed in quotation marks and separated by commas.

<u>Response 1</u>

"ACTB","GAPDH","RPL13A","RPS29","RPLP0","TUBB","UBC","YWHAZ","B2M","HPRT1",
"MS4A1","CD19","CD79A","CD79B","PAX5","BCL2","BCL6","MYC","PIM1","EZH2",
"CREBBP","SOCS1","TNFRSF14","CD22","IRF8","CD3D","CD3E","CD3G","CD4","CD8A",
"CD8B","FOXP3","CTLA4","PDCD1","ICOS","TRAC","TRBC1","TRBC2","NCAM1","KLRD1",
"NKG7","GNLY","FCGR3A","KIR2DL1","KIR3DL1","GZMB","CD14","CD68","CD163","ITGAX",
"CLEC9A","LYZ","FCER1A","CD1C","MARCO","MERTK","TNFRSF8","FUT4","STAT6","JAK2",
"REL","NFKB1","PDPN","FAP","COL1A1","COL1A2","DCN","BGN","THY1","VIM",
"MMP2","MMP9","PECAM1","VWF","ENG","CD34","ICAM1","IL2","IL4","IL6",
"IL7","IL10","IL12A","IL15","IL17A","IL1B","IFNG","TNF","TGFB1","CCL2",
"CXCL9","CXCL10","CXCL11","CXCL13","CCL19","CCL21","CXCL12","CXCL14","CCR7","CXCR5",
"CD28","SLAMF1","PDCD1LG2","CD274","CD80","CD86","CD40","CD40LG","CD70","CD27",
"TNFRSF9","TNFSF9","MKI67","PCNA","CDK1","CDK2","CCND1","CCNE1","CDKN1A","E2F1",

"AKT1","AKT2","PTEN","PIK3CA","MAPK1","MAPK3","MAPK8","MAPK14","NFATC1","JAK1",
"TYK2","STAT1","STAT3","STAT5A","NFKBIA","RELA","BRAF","KRAS","ERK","SRC",
"LDHA","HK2","PKM","GPI","PGK1","PFKP","SLC2A1","SLC16A1","MTOR","AMPK",
"CD44","CDH1","CDH2","SNAI1","TWIST1","ZEB1","ZEB2","FN1","SPARC","VCAM1",
"DNMT1","TET2","KDM6B","KMT2A","HDAC1","HDAC2","SMARCA4","ARID1A","BRD4",
"EP300",
"TP53","RB1","CDKN2A","ATM","BRCA1","BRCA2","PTPN11","ETS1","FOS","JUN",
"CASP3","CASP8","CASP9","BID","BAX","BAK1","MCL1","XIAP","BIRC5","FAS",
"TNFAIP3","CARD11","NFKB2","RELB","IRF4","KMT2D","HIST1H1E","MYD88","CD2","IGHM"

<u>Prompt 2</u>

I am designing a gene panel consisting of 200 genes in the context of lymph node cancer in humans. The goal is to optimize the panel based on the following criteria:

- Cell type specific markers, where the cell types of interest are: B cells, plasma cells, monocyte/macrophage, fibroblasts, endothelial cells, and malignant cells.

- Transcriptional variation, try to capture as much of the transcriptional variation possible.
- Feature diversity, ensuring minimal redundancy among positively correlated features.
- Inclusion of diverse pathways to capture broad biological relevance.
- Representation of ligands and receptors, crucial for cell-cell communication.
- Inclusion of spatially variable genes, capturing spatial heterogeneity in AML.

Please design a gene panel of 200 genes based on these criteria. Provide the gene list in one continuous line, with each gene enclosed in quotation marks and separated by commas.

<u>Response 2</u>

"ACTB","GAPDH","RPL13A","RPS29","RPLP0","TUBB","UBC","YWHAZ","B2M","HPRT1",
"MS4A1","CD19","CD79A","CD79B","PAX5","CD22","CR2","BLK",
"PRDM1","IRF4","CD27","CD38","SDC1","MZB1","XBP1","JCHAIN",
"CD14","FCGR3A","CD68","MARCO","CSF1R","ITGAX","SIRPA","CD86",
"PDPN","FAP","PDGFRA","PDGFRB","DCN","COL1A1","COL1A2","CXCL12","CXCL14","BGN","ELN","MMP2","MMP3","THY1","VIM",
"PECAM1","VWF","ENG","TIE1","TEK","CD34","FLT1","KDR","NOS3","ICAM1",
"BCL2","BCL6","MYC","PIM1","EZH2","CREBBP","SOCS1","TNFRSF14","IRF8","FOXP1","NOTCH1","TP53","RB1","CDKN2A","ATM","BRCA1","BRCA2","PTEN","KRAS","NRAS","BRAF","KIT","FLT3","IDH1","IDH2","NPM1","TET2","DNMT3A","RUNX1","CEBPA","MPO",
"IL2","IL4","IL6","IL7","IL10","IL12A","IL12B","IL15","IL17A","IL1B","IFNG","TNF","TGFB1","CXCL9","CXCL10","CXCL11","CXCL13","CCL19","CCL21","CCL2",
"CTLA4","PDCD1","CD274","PDCD1LG2","CD80","ICOS","CD40","CD40LG","TNFRSF9","TNFSF9",
"MAPK1","MAPK3","MAPK8","MAPK14","AKT1","AKT2","JAK1","JAK2","STAT3","NFKB1",
"KMT2A","KMT2D","HDAC1","HDAC2","SMARCA4","ARID1A",

"FN1","VCAN","LAMA1","LAMC2","VTN","FDCSP","MADCAM1","LYVE1","ACKR4","CCR5","CXCR5","ICAM2","VCAM1","SELE","SELPLG","SELL","TGFBI",
"TNFRSF1A","TNFRSF1B","TNFRSF4","TNFSF4","TNFRSF18","TNFSF18","TNFSF14","CD27","CD70","TNFRSF13B",
"TCF3","LEF1","GATA3","TBX21","RORC","RUNX3","MAF","ETS1","FOS","JUN","CARD11","TCF4","BACH2","E2F2","MDM2","MDM4","ETV6","FOXO1","FOXO3","STAT5A","STAT5B","SMAD2","SMAD3","SMAD4","RAD51","HIF1A","BCL3",
"LDHA","HK2","PKM","GPI","PGK1","PFKP","SLC2A1","SLC16A1","MTOR","PRKAA1"

### Supplementary 3

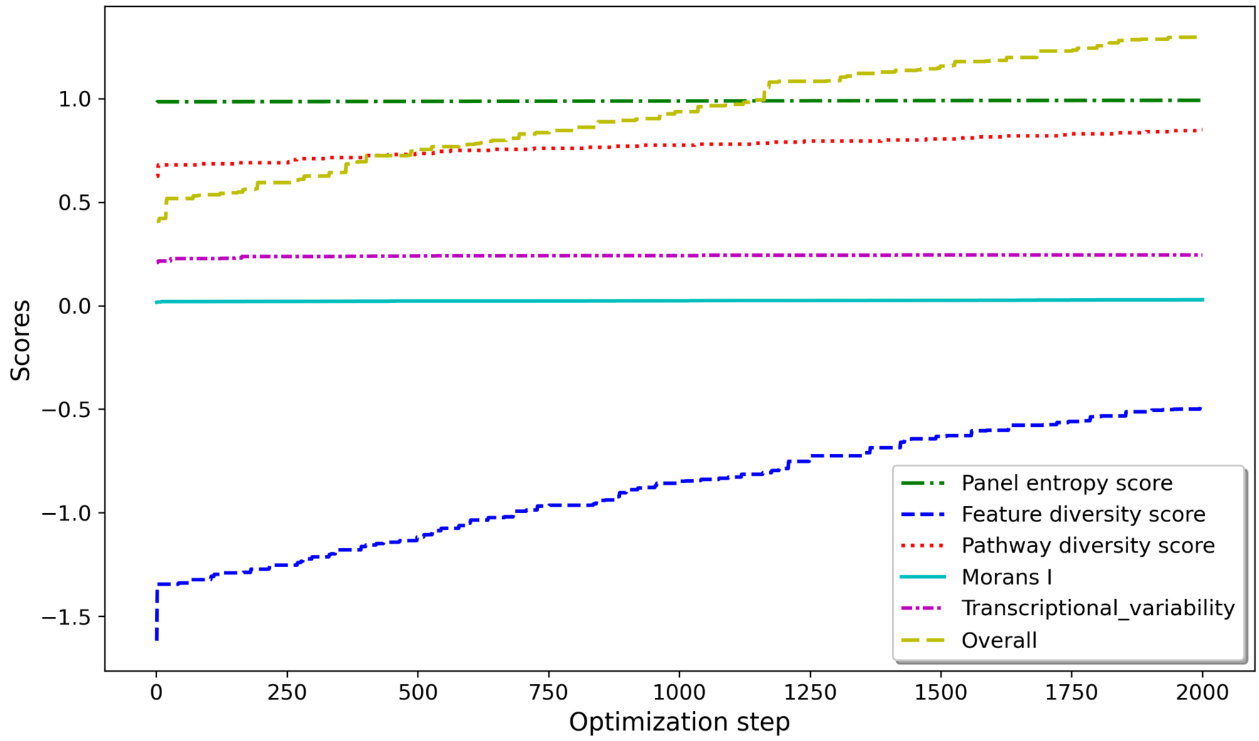


Metric changes over the course of genetic algorithm iterations. The plot shows the progress of five objective functions— Panel entropy score, Feature diversity score, Pathway diversity score, Moran’s I and transcriptional-variability-based—tracked against their overall metric. The y-axis represents the score values, while the x-axis represents the optimization steps. All metrics show steady increases in score throughout the optimization process, indicating improved performance with continued iterations. Some flat lines are only due to the drawing reasons (the range of y-axis) and would show increase when zoomed in (exemplified as Supplementary 4).

### Supplementary 4

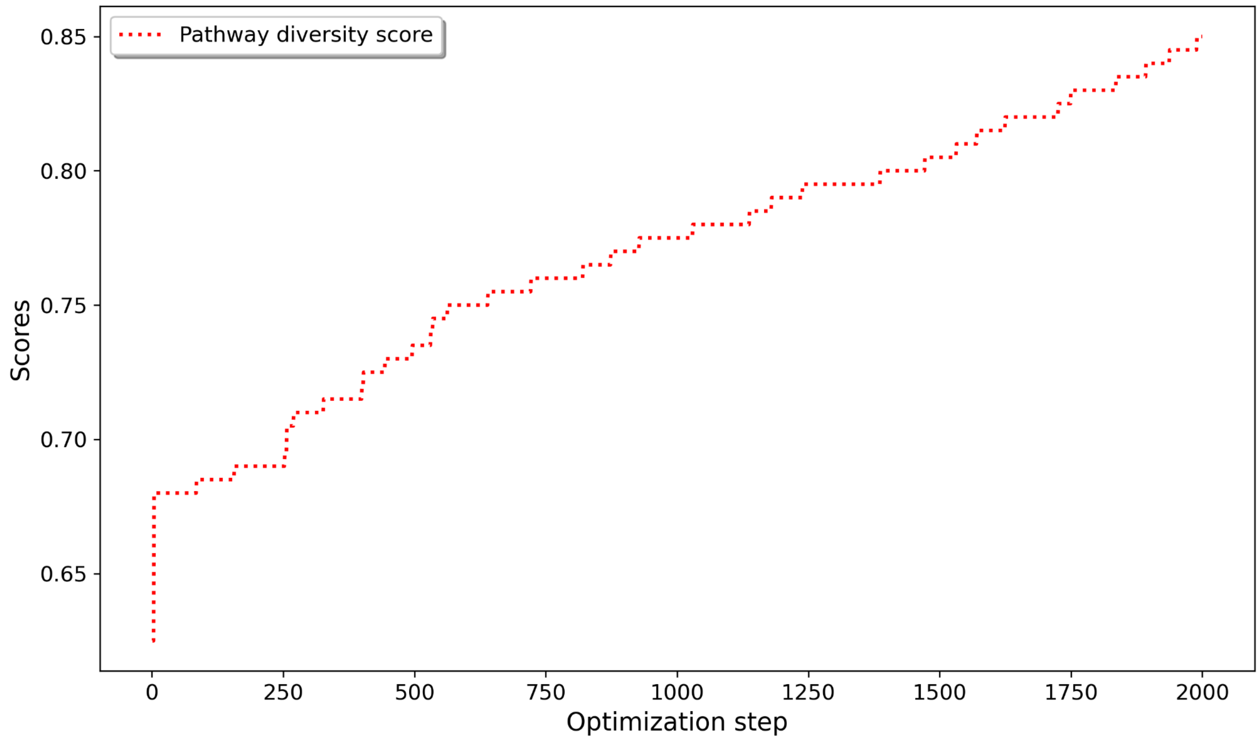


Pathway diversity score changes over the course of genetic algorithm iterations with a zoomed-in perspective mentioned in Supplementary 3.
